# SCAPE-APA: a package for estimating alternative polyadenylation events from scRNA-seq data

**DOI:** 10.1101/2024.03.12.584547

**Authors:** Guangzhao Cheng, Tien Le, Ran Zhou, Lu Cheng

## Abstract

**Summary:** SCAPE is a package we previously developed to estimate alternative polyadenylation events from single cell RNA-seq (scRNA-seq) data, which is composed of ad-hoc python scripts and has speed issues when handling large scRNA-seq data. To suit the needs of analyzing large scRNA-seq datasets, we present SCAPE-APA, which is a re-implementation of SCAPE with substantial changes. We made the following updates to the package (1) we binned similar reads together to accelerate the estimation (2) we re-derived the mixture model to tailor it for binned reads (3) we implemented the inference algorithm using Taichi language for acceleration (4) we re-implemented the untranslated region (UTR) annotation extraction script using the professional package gffutils for better maintenance (5) we wrote a script to detect spurious alternative polyadenylation sites generated due to junction reads and (6) we made a formal python package and uploaded it to the Python Package Index website (Pypi).

**Availability and Implementation:** Scape-apa is freely available at https://github.com/chengl7-lab/scape and can be easily installed using pip.

## Introduction

The 3’ untranslated region (3’UTR) of a messenger RNA (mRNA) molecule is a critical segment located downstream of the coding sequence and plays a pivotal role in gene regulation. Polyadenylation sites (pA sites) are specific locations within the 3’UTR where the mRNA undergoes a process known as polyadenylation, resulting in the addition of a poly(A) tail to a pA site. Alternative polyadenylation (Mitschka and Mayr 2022) refers to the phenomenon where a single gene can possess multiple pA sites within its 3’UTR, leading to the production of distinct mRNA isoforms with varying 3’UTR lengths.

These alternative polyadenylation isoforms can have profound implications for post-transcriptional regulation, as they may influence mRNA localization, stability, and interaction with regulatory factors, thereby impacting the ultimate functional output of the gene. Understanding the mechanisms and consequences of alternative polyadenylation is essential for unraveling the complexities of gene expression and its role in diverse biological processes.

We previously developed a software package SCAPE (Zhou et al. 2022) that can estimate alternative polyadenylation events from single cell RNA-seq (scRNA-seq) data. Since the reads in scRNA-seq are generally not long enough to cover the pA sites, we used a Bayesian mixture model (Fig. 1A) to infer the pA sites by utilizing the insert length distribution. The mixture model contains *k* (*k* > 1) Gaussian components and one uniform component. Each pA site is represented by a Gaussian component, which has three parameters: the mean *μ*, the standard deviation *δ* and the weight *π* . The full SCAPE analysis workflow can be summarized into the following steps: (1) extract 3’UTR annotation from the GTF (General Transfer Format) or GFF (General Feature Format) file (2) map the reads to the reference genome and calculate the relative positions of reads on the corresponding 3’UTR regions (3) perform parameter inference for each 3’UTR region and output the result (4) perform downstream tasks such as differential pA site usage analysis.

**Figure 1.**
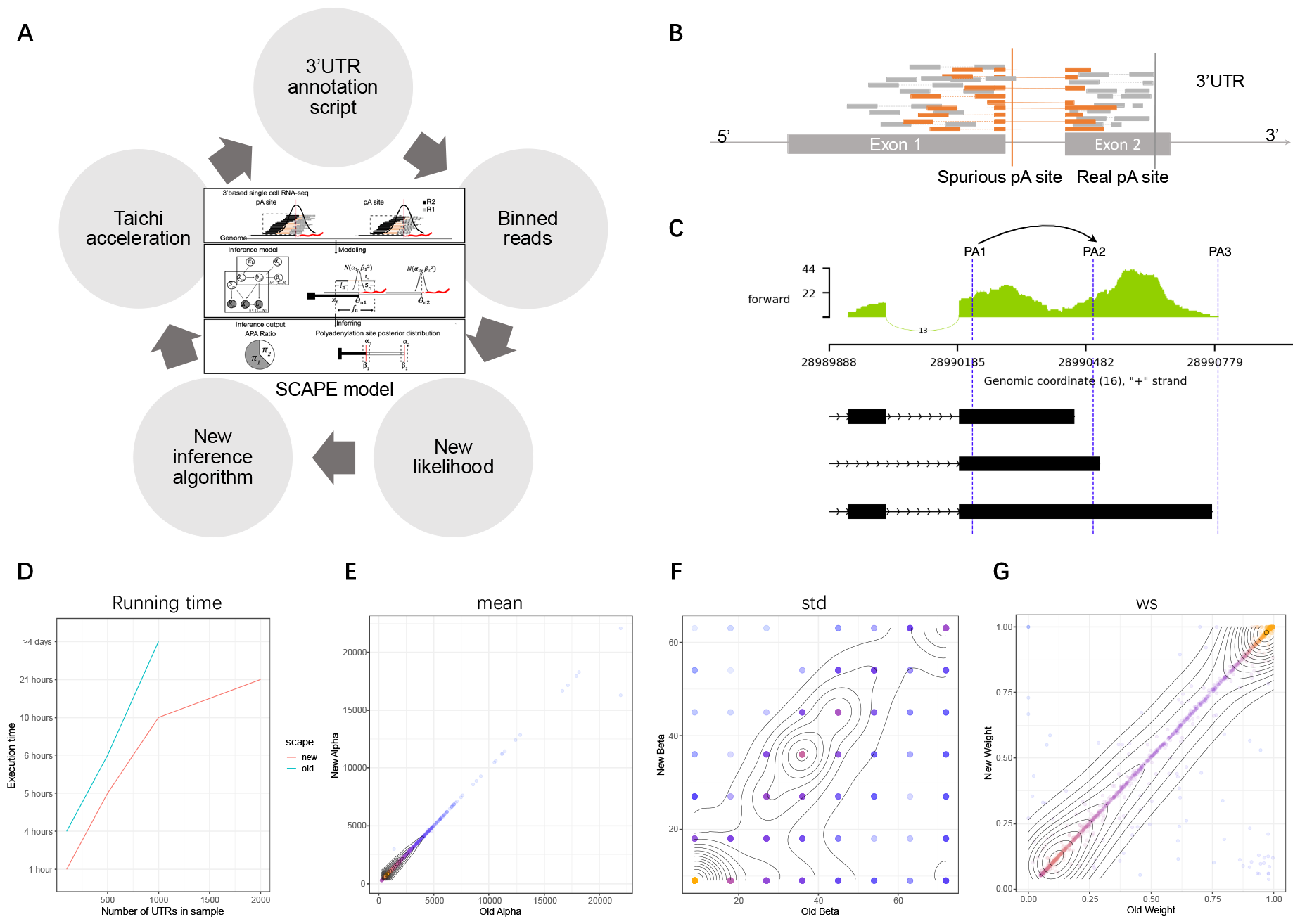
(**A**) Original SCAPE model and improvements of SCAPE-APA. (**B**) Schematic illustration of a spurious pA site due to splicing of 3’UTR. (**C**) A real spurious pA site case. PA1 is the spurious pA site and should be merged into PA2. (**D**) Comparison of running time on the same computer across different datasets. (**E-G**) Comparison of estimated mean, std and weight of estimated pA sites. The curves are the density contour of data points.

### Improvements

We have made significant updates to the previous implementation, as highlighted in Fig. 1A. The biggest challenge in real data analysis is that the read number can be very large (*n* > 200,000) for some 3’UTRs. This has made the original inference algorithm extremely slow. As the 3’UTR is relatively short (∼1500bp), there are lots of redundant reads. To solve this problem, we binned reads in the vicinity together, i.e. reads with similar *x, l, r* values in the following likelihood.

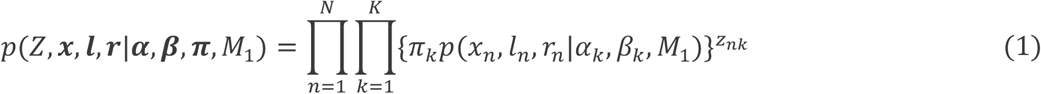

Note that the bin is 3-dimensional. Let us denote the *n*th bin by the unique combination  *x*_*n*_, *l*_*n*_, *r*_*n*_) and its count by *c*_*n*_, where the bin sizes are set to 5bp, 10bp, 10bp, respectively. We can rewrite the joint likelihood of the mixture model as follows:

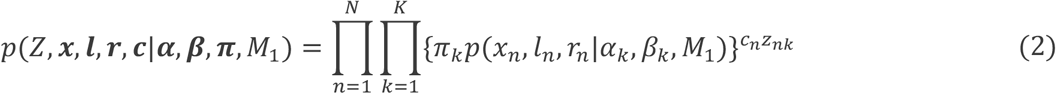

It can be seen that we only need to calculate the likelihood once for all *c*_*n*_ reads in the *n*th bin in the new likelihood, while this has to be repeated *c*_*n*_ times in the original likelihood. As the likelihood has changed, the inference algorithm has to be changed accordingly. Detailed mathematical derivations of the inference algorithm for the new likelihood are provided in Supplementary Note 1. To further accelerate the inference, we use Taichi language (Hu et al. 2019) to re-implement the inference algorithm, where Taichi uses the just-in-time (JIT) compilation to accelerate python code. Since the inference algorithm is serial, the computation times are similar no matter if GPU is used or not.

Beside the new likelihood and inference algorithm, we re-implemented and added new bioinformatic functionalities to streamline the workflow. We re-wrote the python script to extract 3’UTR annotations from GTF or GFF files, based on the package “gffutils” (https://github.com/daler/gffutils), which was easier to maintain compared with the previous ad hoc implementation. We added a post-processing script to handle spurious pA sites due to junction reads. As shown in Fig. 1B, some 3’UTR may contain multiple exons/fragments, while the SCAPE model assumes 3’UTR to be a consecutive region on the genome, i.e. we take the start of the first exon/fragment and the end of the last exon/fragment as the full 3’UTR region. Due to this multi-segment property of the given 3’UTR in Fig. 1B, two pA sites were estimated from the data, where the orange pA site was the spurious pA site. To resolve this, we merged a pA site with the next downstream pA site if there were more than 40% junction reads in all reads assigned to this pA site. Fig. 1C shows a spurious pA site case in the real data, where the first estimated pA site should be merged with the second. Finally, we made a python package “scape-apa” and uploaded it to the Python Package Index (PyPI) to facilitate the installation.

We compared the speed and inference results of the old and new SCAPE software by sampling 100, 500, 1000, 2000 3’UTRs from a public scRNA-seq data (Notaras et al. 2022). Fig. 1D shows the running times for analyses of these datasets on a CPU-only computer (Dell PowerEdge C4130), with the maximum memory set to 10GB. It can be seen that the speedup is 9-fold for the dataset with 1000 3’UTRs. Fig. 1E-G show the estimated mean, std and weights given by the old and new SCAPE package of the dataset with 1000 3’UTR. We fixed the number of pA sites in each UTR according to the results of the new package. It can be seen that the majority of the data points are scattered around the *y*=*x* line for the mean and weight, while larger variation is observed for the standard deviation. These differences are due to the binning operation, but overall the comparisons suggest a highly consistent result between the old and new packages.

In summary, we have revised the statistical model for binned reads and utilized Taichi to accelerate the parameter inference of SCAPE, as well as developed useful python scripts to process GTF files and handle spurious pA sites.

## Availability

The SCAPE-APA implementation, benchmark datasets and scripts are provided in https://github.com/chengl7-lab/scape.

## Competing interests

The authors declare no competing interests.

## Funding

This work was supported by the Research Council of Finland [grant number 335858, 358086 to G. C., T. L. and L. C.].

## Author contribution

G. C. developed the inference algorithm for binned reads and implemented the algorithm in Taichi. T. L. developed the python scripts for parsing GTF file and removing spurious pA sites and ran the benchmark experiments. R. Z. provided advice about 3’UTR annotation extraction. L. C. supervised the whole project.

## Notes

### Competing Interest Statement

The authors have declared no competing interest.

## References

Hu, Yuanming, et al. (2019), ‘Taichi: a language for high-performance computation on spatially sparse data structures’, ACM Transactions on Graphics, 38 (6), 1–16.

Mitschka, S. and Mayr, C. (2022), ‘Context-specific regulation and function of mRNA alternative polyadenylation’, Nat Rev Mol Cell Biol, 23 (12), 779–96.

Notaras, M., et al. (2022), ‘Schizophrenia is defined by cell-specific neuropathology and multiple neurodevelopmental mechanisms in patient-derived cerebral organoids’, Mol Psychiatry, 27 (3), 1416–34.

Zhou, R., et al. (2022), ‘SCAPE: a mixture model revealing single-cell polyadenylation diversity and cellular dynamics during cell differentiation and reprogramming’, Nucleic Acids Res, 50 (11), e66.

