## Supplementary Note 1 for "SCAPE-APA: a package for estimating alternative polyadenylation events from scRNA-seq data"

### Statistial model of SCAPE-APA

(Supplementary Note 1)

Guangzhao Cheng, Lu Cheng

February 29, 2024

#### 1 Background

SCAPE stands for Single Cell Alternative Polyadenylation using Expectation-maximization. This document tries to establish a mixture model to estimate alternative polyadenylation (APA) events from scRNA-seq datasets. APA is a common type of splicing in the 3'-untranslated region (3'-UTR) near the ployA part of mRNA, where 3'-UTR is cleaved at different polyadenylation (pA) sites and polyA tails are appended. The mixture model aims to identify the pA sites and quantify the proportions of different isoforms from polyA-captured scRNA-seq such as 10X.

10X sequencing is a single cell sequencing technique that captures mRNA from a single cell. The protocol first capture the polyA tails using a 30bp polyT (followed by UMI and barcode) on a bead, then break the captured mRNA using sonification. After that cDNA is generated and followed by PCR. DNA fragments from different cells are pooled together, which is subject to Illumina pair-end sequencing. Illumina pair-end sequencing in general only pick up DNA fragments around 300bp, with 50bp standard deviation. The pair-end read is around 150bp.

Note that the 30bp polyT may bind to any part of polyA tails (20-150bp, [1]) in mRNA capture. In principle, we could directly measure the PA sites since some reads can directly cover the joint part between polyA and UTR, which we term as **junction reads**. However, the sequencing quality deteriorate dramatically after sequencing long stretch of polyA due to technical reasons. Therefore, the non-polyA part of R1 of many reads cannot be successfully mapped back to 3'-UTR on the genome.

The proportion of junction reads is very low (Supplementary Note Figure 2), which provide limited information about the location of pA sites and the proportions of isoforms. Given the length distribution of DNA fragments in Illumina sequencing, as well as the length distribution of polyA tail, we could infer the positions of pA sites and their proportions of the corresponding isoforms using statistical modelling. In order to answer this question, we propose a mixture model in the sequel.

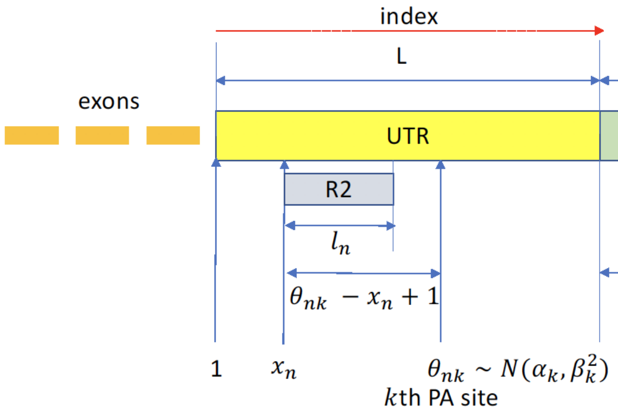


Supplementary Note Figure 1: APA isofrom model. $\theta_{nk}$ is the $k$th pA (cleavage) site where the UTR and polyA connect on $n$th DNA fragment (pair-end read). $\alpha_k$ and $\beta_k$ are the mean and standard deviation of the $k$th pA site. $x_n$ is the start position relative to the start of UTR, for the DNA fragment corresponding to the $n$th pair-end reads.

### 2 Model

Based on the existing knowledge, we propose a mixture model as shown in Supplementary Note Figure 1. The yellow UTR refers to the full UTR on the genome, whose length is $L$. We assume the maximum length of the green polyA tail is 150bp, according to a previously published data [1]. For the $k$th ($1 \leq k \leq K$) isoform and $n$th ($1 \leq n \leq N$) pair-end read, we assume the position of the pA site is $\theta_{nk}$, i.e. isoform $k$ will connect with polyA tails at

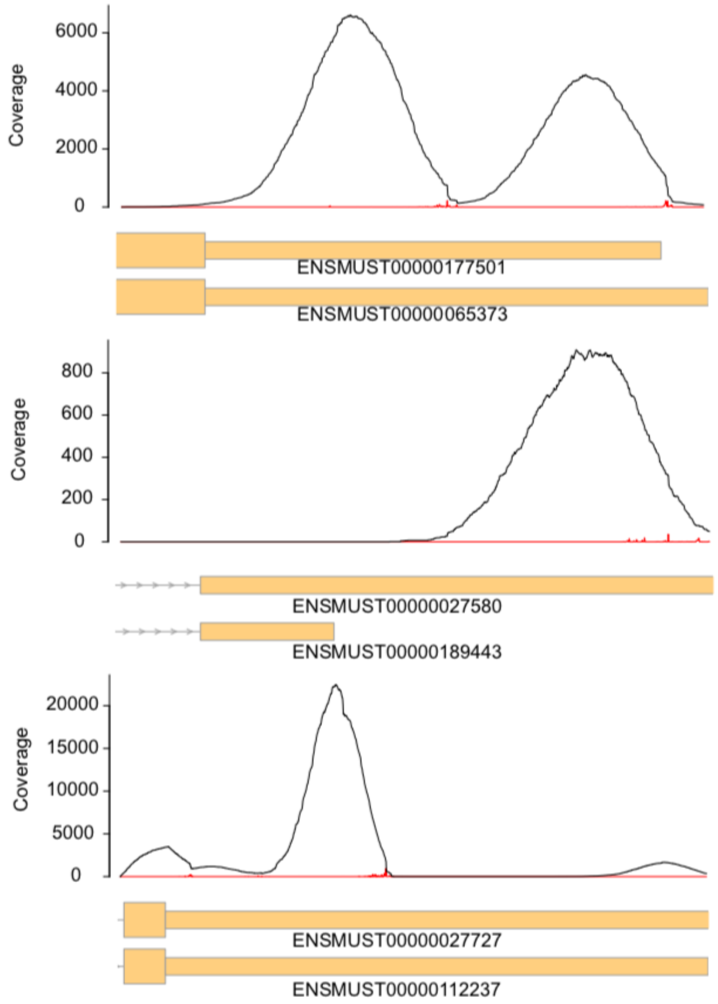


Supplementary Note Figure 2: Junction reads of three example genes from the 10X mouse bone marrow dataset. The black curve is the coverage of scRNA-Seq reads mapped to UTR. The red lines represented the coverage of pA sites (only 1bp) given by junction reads. pA site annotations from NCBI database are provided beneath the x-axis. It can be seen that the amount of junction reads is very small and the pA sites exhibit a certain degree of fluctuation.

position $\theta_{nk}$. We assume for the $n$th pair-end read, the end on the UTR part is called by $R2$ and the end on the polyA part is called by $R1$, the lengths

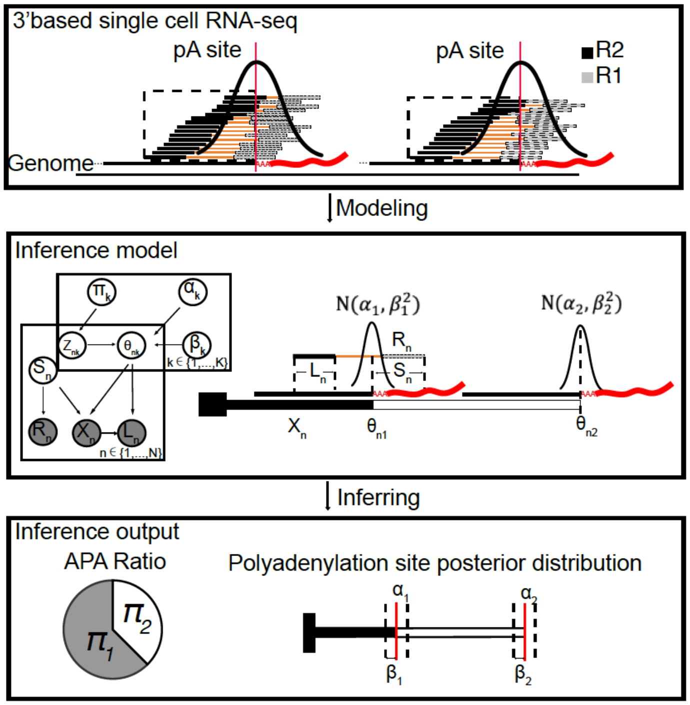


Supplementary Note Figure 3: SCAPE model. Pair-end reads are first mapped to the genome and reads land on the UTR regions are extracted. Bioinformatic analysis is performed to extract relevant information from the reads, including $l_n$, $r_n$ and $x_n$. With the preprocessed data fed into the model, it will output estimated parameters of the locations of pA sites ($\alpha_k$, $\beta_k$) and their proportions ($\pi_k$).

of which are denoted by $l_n$ and $r_n$, respectively. The break position on the UTR part of $n$th pair-end read is denoted by $x_n$, i.e. R2 maps to position $x_n$. Thus, the length of the UTR part in the fragment is $\theta_{nk} - x_n + 1$. We denote the maximum length of polyA by $LA = 150$ and the length of the polyA part in the fragment by $s_n$, which follows an empirical distribution estimated from fully mapped junctions reads of the 10X mouse bone marrow dataset. By investigating the limited junction reads in our data, a pA site usually exhibits a certain amount of fluctuation, as shown in Supplementary Note Figure 2. Therefore, we assume the $k$th pA site $\theta_{nk}$ is not fixed, but rather sampled from a Gaussian distribution $N(\alpha_k, \beta_k^2)$, where $\alpha_k$ is the mean position and $\beta_k$ is the standard deviation.

Note that $x_n, l_n, r_n$ are observed variables, $\theta_{nk}, s_n$ are hidden variables that needs to be marginalized out. The proportions of different isoforms $\boldsymbol{\pi}$, parameters of pA site ($\boldsymbol{\alpha}$, $\boldsymbol{\beta}$) are the parameters to be estimated.

Also, there exists reads that cannot be well explained by the APA isofrom model. We treat these reads as random noise, which are uniformly sampled from the UTR and polyA part.

#### 2.1 Data generating process

The whole workflow of SCAPE is provided in Supplementary Note Figure3. The generative model is given in the central panel. Detailed data generating process is given as follows

1. Specify $K+1$ components and the corresponding weights $\boldsymbol{\pi} = (\pi_0, \pi_1, \ldots, \pi_K)$. Note we have $\sum_{k=0}^{K} \pi_k = 1$ and $\pi_k > 0$. $k = 0$ indicates the noise components and $k > 0$ refers to APA isoform component.

2. For isoform $k$, sepcify $\alpha_k$ and $\beta_k$. Note we have $\alpha_k \in \{1, 2, \ldots, L\}$ and $\beta_k > 0$.

3. For read pair $n$, sample a component label $k$ from $Cat(\pi_0, \pi_1, \ldots, \pi_K)$. If $k = 0$, then sample the reads following steps in 5; otherwise sample the reads following steps in 4.

4. Sample reads from APA isofrom model.

   (a) Set the indicator variable $z_{nk} = 1$ for component $k$ and $z_{nj} = 0$ for $j \neq k$.

   (b) Sample $\theta_{nk}$ from a Gaussian distribution $\theta_{nk} \sim N(\alpha_k, \beta_k^2)$.

   (c) Generate the length of polyA part in the fragment $s_n$ from uniform distribution Unif(20, 150), or from an empirical distribution (Categorical). Then generate the length $r_n$ from uniform distribution Unif(10,$s_n$).

   (d) Generate fragment length from $f_n \sim N(\mu_f, \sigma_f^2)$, where $\mu_f = 300$ and $\sigma_f = 30$. The break position on the UTR part of the fragment $x_n$ is given by $\theta_{nk} - (f_n - s_n) + 1$.

   (e) Generate the length $l_n$ from uniform distribution Unif(10,$\theta_{nk} - x_n + 1$).

(f) The outputs are $x_n$, $l_n$ and $r_n$. $\theta_{nk}$, $s_n$ , $f_n$ and $z_{nk}$ are hidden variables.

5. Sample reads from noise model.

   (a) Set the indicator variable $z_{nk} = 1$ for component $k$ and $z_{nj} = 0$ for $j \neq k$.

   (b) Generate the length $r_n$ from uniform distribution Unif(10,150).

   (c) Sample the break position on the UTR part $x_n$ from Unif(1,$L$).

   (d) Sample the length $l_n$ from Unif(1,$L - x_n$).

To improve computation efficiency, we bin similar data points into bins based on $\boldsymbol{x}$, $\boldsymbol{l}$, $\boldsymbol{r}$. After the binning operation, the data for the $n$th bin become $x_n$, $l_n$, $r_n$ and $c_n$, where $c_n$ is the total number of data points in $n$th bin and we use $\boldsymbol{c}$ to denote counts for all bins. $x_n$, $l_n$ and $r_n$ are the mean of data points fall into $n$th bin.

Next we first provide the mathematical definitions of the APA isoform model, noise model separately; then introduce the full mixture model.

#### 2.2 APA isofrom model

First we look at the APA isofrom model $M_1$. Given the isoform weights $\boldsymbol{\pi}$ and pA site parameters $\boldsymbol{\alpha}$ and $\boldsymbol{\beta}$. The joint likelihood is given by

$$p(Z, \boldsymbol{x}, \boldsymbol{l}, \boldsymbol{r}, \boldsymbol{c}|\boldsymbol{\alpha}, \boldsymbol{\beta}, \boldsymbol{\pi}, M_1) = \prod_{n=1}^{N}\prod_{k=1}^{K}\{\pi_k p(x_n, l_n, r_n|\alpha_k, \beta_k, M_1)\}^{c_n z_{nk}}, \tag{1}$$

where

$$p(x_n, l_n, r_n|\alpha_k, \beta_k, M_1) = \sum_{\theta_{nk}} p(x_n, l_n, r_n, \theta_{nk}|\alpha_k, \beta_k) \tag{2}$$

$$= \sum_{\theta_{nk}} p(x_n, l_n, r_n|\theta_{nk})p(\theta_{nk}|\alpha_k, \beta_k) \tag{3}$$

$$= \sum_{\theta_{nk}} p(x_n, l_n, r_n|\theta_{nk})N(\theta_{nk}|\alpha_k, \beta_k^2) \tag{4}$$

The first likelihood term is given by

$$p(x_n, l_n, r_n|\theta_{nk}) = \sum_{s_n} p(x_n, l_n, r_n, s_n|\theta_{nk}) \tag{5}$$

$$= \sum_{s_n} p(l_n|x_n, \theta_{nk})p(x_n|s_n, \theta_{nk})p(r_n|s_n)p(s_n) \tag{6}$$

$$= \sum_{s_n} f(x_n, l_n, r_n, s_n, \theta_{nk}), \tag{7}$$

where we have defined $f(\cdot)$ to simplify notation by

$$f(x_n, l_n, r_n, s_n, \theta_{nk}) = p(l_n|x_n, \theta_{nk})p(x_n|s_n, \theta_{nk})p(r_n|s_n)p(s_n) \tag{8}$$

The distributions of the variables are given as follows.

$$p(s_n) = Unif(20, 150), \tag{9}$$

where we assume minimum polyA length is 20bp. Also it could be empirical distribution (categorical) estimated from existing datasets.

$$p(r_n|s_n) = \begin{cases} \frac{1}{s_n}, & \text{if } 1 \leq r_n \leq s_n \\ 0, & \text{otherwise} \end{cases}$$

Note that the capturing polyT can bind to any position on the polyA tail, so the denominator is $s_n$. For reads of the polyA end (R1 in Supplementary Note Figure 1) that covers pA sites, i.e. $s_n$ is known, we set $p(s_n)$ and $p(r_n|s_n)$ to 1, since the length $r_n$ is fully determined. Note that the non-polyA part of these reads may not match UTR.

$$p(x_n|s_n, \theta_{nk}) = N(\theta_{nk} + s_n + 1 - \mu_f, \sigma_f^2), \tag{10}$$

where $\mu_f$ and $\sigma_f$ are expected mean and standard deviation of fragment length. Note that the $n$th fragment length is $\theta_{nk} - x_n + 1 + s_n \sim N(\mu_f, \sigma_f^2)$, and $x_n$, $s_n$, $\theta_{nk}$ are known. Therefore, we have $x_n|s_n, \theta_{nk} \sim N(\theta_{nk} + s_n + 1 - \mu_f, \sigma_f^2)$.

$$p(l_n|x_n, \theta_{nk}) = \begin{cases} \frac{1}{\theta_{nk} - x_n + 1}, & \text{if } 1 \leq l_n \leq \theta_{nk} - x_n + 1 \\ 0, & \text{otherwise} \end{cases}$$

. When deriving $l_n$ for junction reads (R2 in Supplementary Note Figure1), we trim away the polyA parts, which leads to low probabilities for these reads. To compensate this, we set the probability of $p(x_n|s_n, \theta_{nk})$ to 1. This treatment will give such reads more weights in inferring the pA sites.

#### 2.3 Noise model

To account for random noise, we use a noise model $M_0$ to model random reads that do not support pA sites. For $n$th read, the likelihood of the noise model is given by

$$
\begin{aligned}
p(x_n, l_n, r_n, \alpha_0, \beta_0 | M_0) &= p(x_n, l_n, r_n | M_0) &(11)\\
&= p(x_n | M_0) p(l_n | s_n, M_0) p(r_n | s_n, M_0) &(12)\\
&= \frac{1}{L} \cdot \frac{1}{L} \cdot \frac{1}{LA}, &(13)
\end{aligned}
$$

where $\alpha_0$ and $\beta_0$ are not used but kept for notation simplicity; $L$ is the length of the UTR part; $LA$ is the maximum length of polyA tails and

$$
\begin{aligned}
p(x_n | M_0) &= \frac{1}{L} &(14)\\
p(l_n | s_n, M_0) &= \frac{1}{L} &(15)\\
p(r_n | s_n, M_0) &= \frac{1}{LA} &(16)
\end{aligned}
$$

#### 2.4 Full Model

The likelihood of the full model is given by

$$
\begin{aligned}
&p(Z, \boldsymbol{x}, \boldsymbol{l}, \boldsymbol{r}, \boldsymbol{c} | \boldsymbol{\alpha}, \boldsymbol{\beta}, \boldsymbol{\pi}) &(17)\\
&= \prod_{n=1}^{N} \left\{ \prod_{k=1}^{K} \{\pi_k p(x_n, l_n, r_n | \alpha_k, \beta_k, M_1)\}^{c_n z_{nk}} \{\pi_0 p(x_n, l_n, r_n | \alpha_0, \beta_0, M_0)\}^{c_n z_{n0}} \right\} &(18)
\end{aligned}
$$

The log-likelihood of the full model is given by

$$
\log p(Z, \boldsymbol{x}, \boldsymbol{l}, \boldsymbol{r}, \boldsymbol{c} | \boldsymbol{\alpha}, \boldsymbol{\beta}, \boldsymbol{\pi}) = \sum_{n=1}^{N} \sum_{k=0}^{K} c_n z_{nk} \{\log \pi_k + \log p(x_n, l_n, r_n | \alpha_k, \beta_k, M_{I(k)})\}, \quad (19)
$$

where $I(k) = 1$ if $k \geq 1$ and $I(k) = 0$ if $k = 0$.

### 3 Inference

Here we provide the detailed derivations for the parameter inferences using expectation maximization (EM) algorithm.

The log joint likelihood is given by

$$\log p(Z, \boldsymbol{x}, \boldsymbol{l}, \boldsymbol{r}, \boldsymbol{c}|\boldsymbol{\alpha}, \boldsymbol{\beta}, \boldsymbol{\pi}) \tag{20}$$

$$= \sum_{n=1}^{N} c_n \left\{ \sum_{k=1}^{K} z_{nk}\{\log \pi_k + \log p(x_n, l_n, r_n|\alpha_k, \beta_k, M_1)\} + z_{n0}\eta_0 \right\} \tag{21}$$

$$= \sum_{n=1}^{N} c_n \left\{ \sum_{k=1}^{K} z_{nk}\{\log \pi_k + \log \sum_{\theta_{nk}} p(x_n, l_n, r_n|\theta_{nk})p(\theta_{nk}|\alpha_k, \beta_k)\} + z_{n0}\eta_0 \right\} \tag{22}$$

$$= \sum_{n=1}^{N} c_n \left\{ \sum_{k=1}^{K} z_{nk}\{\log \pi_k + \log \sum_{\theta_{nk}} p(x_n, l_n, r_n|\theta_{nk})N(\theta_{nk}|\alpha_k, \beta_k^2)\} + z_{n0}\eta_0 \right\}, \tag{23}$$

$$= \sum_{n=1}^{N} c_n \left\{ \sum_{k=1}^{K} z_{nk}\{\log \pi_k + \log \sum_{\theta_{nk}} \sum_{s_n} f(x_n, l_n, r_n, s_n, \theta_{nk})N(\theta_{nk}|\alpha_k, \beta_k^2)\} + z_{n0}\eta_0 \right\}, \tag{24}$$

where $\eta_0 = \log \pi_0 + \log p(x_n, l_n, r_n|\alpha_0, \beta_0, M_0)$. Note that $\theta_{nk}$ and $s_n$ are auxiliary variables, which are integrated out in the calculation of the likelihood.

#### 3.1 E-step

In E-step, we try to estimate the probabilities of read assignment $Z$ given everything else. The posterior probability of $Z$ is given by

$$p(Z|\boldsymbol{x}, \boldsymbol{l}, \boldsymbol{r}, \boldsymbol{c}, \boldsymbol{\alpha}, \boldsymbol{\beta}, \boldsymbol{\pi}) \propto p(Z, \boldsymbol{x}, \boldsymbol{l}, \boldsymbol{r}, \boldsymbol{c}|\boldsymbol{\alpha}, \boldsymbol{\beta}, \boldsymbol{\pi}) \tag{25}$$

$$= \prod_{n=1}^{N} \left\{ \prod_{k=1}^{K} \{\pi_k p(x_n, l_n, r_n|\alpha_k, \beta_k, M_1)\}^{c_n z_{nk}} \{\pi_0 p(x_n, l_n, r_n|\alpha_0, \beta_0, M_0)\}^{c_n z_{n0}} \right\} \tag{26}$$

Given $\boldsymbol{x}, \boldsymbol{l}, \boldsymbol{r}, \boldsymbol{c}, \boldsymbol{\pi}, \boldsymbol{\alpha}, \boldsymbol{\beta}$, the bases in exponential terms can be treated as constants w.r.t. $Z$. The read assignment $z_{nk}$ is a binary value and follows a

Bernoulli distribution. The expectation of $z_{nk}$ is given by

$$\begin{aligned} E(z_{nk}) &= 1 \cdot p(z_{nk}=1) + 0 \cdot p(z_{nk}=0) && (27)\\ &= p(z_{nk}=1) && (28)\\ &\propto \left[\pi_k p(x_n, l_n, r_n | \alpha_k, \beta_k, M_{I(k)})\right]^{c_n}, && (29) \end{aligned}$$

where $I(k) = 1$ if $k \geq 1$ and $I(k) = 0$ if $k = 0$. Therefore, we have

$$E(z_{nk}) = \frac{\left[\pi_k p(x_n, l_n, r_n | \alpha_k, \beta_k, M_{I(k)})\right]^{c_n}}{\sum_{j=0}^{K} \left[\pi_j p(x_n, l_n, r_n | \alpha_k, \beta_k, M_{I(j)})\right]^{c_n}} = \gamma_{nk}, \tag{30}$$

where we have used $\gamma_{nk}$ to denote the expectation in the sequel.

#### 3.2 M-step

In this step, we want to maximize the log likelihood given $Z$ by varying the values of $\boldsymbol{\pi}$, $\boldsymbol{\alpha}$ and $\boldsymbol{\beta}$.

First we try to maximize the log likelihood w.r.t. $\pi_k$, which is constrained by $\sum_{k=0}^{K} \pi_k = 1$. Thus, we introduce the following Lagrange multiplier

$$g(\pi_k) = \log p(Z, \boldsymbol{x}, \boldsymbol{l}, \boldsymbol{r}, \boldsymbol{c} | \boldsymbol{\alpha}, \boldsymbol{\beta}, \boldsymbol{\pi}) + \lambda(1 - \sum_{k=0}^{K} \pi_k) \tag{31}$$

$$= \sum_{n=1}^{N} \sum_{k=0}^{K} c_n z_{nk} \Big\{ \log \pi_k + \log p(x_n, l_n, r_n | \alpha_k, \beta_k, M_{I(k)}) + \lambda(1 - \sum_{k=0}^{K} \pi_k) \Big\} \tag{32}$$

The partial derivative of $g(\pi_k)$ w.r.t. $\pi_k$ is given by

$$\frac{\partial g}{\partial \pi_k} = \sum_{n=1}^{N} c_n \gamma_{nk} \frac{1}{\pi_k} - \lambda \tag{33}$$

By setting $\frac{\partial g}{\partial \pi_k} = 0$, we have $\lambda \pi_k = \sum_{n=1}^{N} c_n \gamma_{nk}$. Therefore, we have

$$\lambda = \lambda \sum_{k=1}^{K} \pi_k = \sum_{k=1}^{K} \lambda \pi_k = \sum_{k=1}^{K} \sum_{n=1}^{N} c_n \gamma_{nk} = \sum_{n=1}^{N} c_n \tag{34}$$

and

$$\pi_k = \frac{1}{\lambda}\sum_{n=1}^{N} c_n \gamma_{nk} = \frac{\sum_{n=1}^{N} c_n \gamma_{nk}}{\sum_{n=1}^{N} c_n} \tag{35}$$

Next we try to maximize the log likelihood w.r.t. $\alpha_k$ and $\beta_k$, but since many terms depend on $\alpha_k$ and $\beta_k$ in a complex way, it is not easy to derive the partial derivatives and analytic solutions of $\alpha_k$ and $\beta_k$. Therefore we try to optimize $\alpha_k$ and $\beta_k$ numerically. We enumerate $\theta_{nk}$ and infer $\alpha_k$ and $\beta_k$ through it. Since $\theta_{nk}$ ranges from 1 to $L$, we could enumerate all its possible values and calculate the corresponding log likelihood. Then we can enumerate reasonable values of $\alpha_k$ $(1 \leq \alpha_k \leq L)$ and $\beta_k$ $(10 \leq \beta_k \leq 70)$. We then choose the value that maximize the log likelihood. In principle we need to enumerate all values, but in practice we enumerate $\theta_{nk}$ every 10 bases.

#### 3.3 Convergence

In order to check the convergence, we provide the lower bound $L(\boldsymbol{\theta})$ as follows

$$\begin{aligned}
L(\boldsymbol{\theta}) &= \sum_{Z} p(Z|\boldsymbol{x}, \boldsymbol{l}, \boldsymbol{r}, \boldsymbol{c}, \boldsymbol{\alpha}^{old}, \boldsymbol{\beta}^{old}, \boldsymbol{\pi}^{old}) \log p(Z, \boldsymbol{x}, \boldsymbol{l}, \boldsymbol{r}, \boldsymbol{c}|\boldsymbol{\alpha}^{old}, \boldsymbol{\beta}^{old}, \boldsymbol{\pi}^{old}) \\
&- \sum_{Z} p(Z|\boldsymbol{x}, \boldsymbol{l}, \boldsymbol{r}, \boldsymbol{c}, \boldsymbol{\alpha}^{old}, \boldsymbol{\beta}^{old}, \boldsymbol{\pi}^{old}) \log p(Z|\boldsymbol{x}, \boldsymbol{l}, \boldsymbol{r}, \boldsymbol{c}, \boldsymbol{\alpha}^{old}, \boldsymbol{\beta}^{old}, \boldsymbol{\pi}^{old}) \\
&= E_Z[\log p(Z, \boldsymbol{x}, \boldsymbol{l}, \boldsymbol{r}, \boldsymbol{c}|\boldsymbol{\alpha}^{old}, \boldsymbol{\beta}^{old}, \boldsymbol{\pi}^{old})] \\
&- \sum_{Z} p(Z|\boldsymbol{x}, \boldsymbol{l}, \boldsymbol{r}, \boldsymbol{c}, \boldsymbol{\alpha}^{old}, \boldsymbol{\beta}^{old}, \boldsymbol{\pi}^{old}) \log p(Z|\boldsymbol{x}, \boldsymbol{l}, \boldsymbol{r}, \boldsymbol{c}, \boldsymbol{\alpha}^{old}, \boldsymbol{\beta}^{old}, \boldsymbol{\pi}^{old}),
\end{aligned}$$

where the expected log-likelihood is given by

$$\begin{aligned}
&E_Z[p(Z, \boldsymbol{x}, \boldsymbol{l}, \boldsymbol{r}, \boldsymbol{c}|\boldsymbol{\alpha}^{old}, \boldsymbol{\beta}^{old}, \boldsymbol{\pi}^{old})] \\
&= \sum_{n=1}^{N}\sum_{k=0}^{K} c_n E(z_{nk}) \left\{ \log \pi_k + \log p(x_n, l_n, r_n|\alpha_k, \beta_k, M_{I(k)}) \right\}
\end{aligned}$$

and the entropy of $Z$ (second term) is given by

$$\sum_{Z} p(Z|\boldsymbol{x}, \boldsymbol{l}, \boldsymbol{r}, \boldsymbol{c}, \boldsymbol{\alpha}^{old}, \boldsymbol{\beta}^{old}, \boldsymbol{\pi}^{old}) \log p(Z|\boldsymbol{x}, \boldsymbol{l}, \boldsymbol{r}, \boldsymbol{c}, \boldsymbol{\alpha}^{old}, \boldsymbol{\beta}^{old}, \boldsymbol{\pi}^{old}) \tag{36}$$

$$= \sum_{n=1}^{N}\sum_{k=1}^{K} c_n \gamma(z_{nk}) \log \gamma(z_{nk}) \tag{37}$$

To compare different models, we use the Bayesian Information Criterion (BIC) given by

$$BIC = -2 \log p(Z, \boldsymbol{x}, \boldsymbol{l}, \boldsymbol{r}, \boldsymbol{c} | \hat{\boldsymbol{\alpha}}, \hat{\boldsymbol{\beta}}, \hat{\boldsymbol{\pi}}) + (3K + 1) \log(N), \tag{38}$$

where $\hat{\boldsymbol{\alpha}}$ , $\hat{\boldsymbol{\beta}}$ and $\hat{\boldsymbol{\pi}}$ are the maximum likelihood estimates. For each APA component, there are one weight parameter $\pi_k$ and two pA site parameters $\alpha_k$, $\beta_k$ for APA component and one parameter $\pi_0$ for the noise component. In total, there are $3K + 1$ free parameters.

##### 3.4 Practical inference

To reduce the computation load, we enumerate $s_n$ from it minimum value (20bp) to its maximum value (150bp) by a 10bp step in the M-step calculation. If some reads cover pA sites, then we will directly extract the hidden variable $s$ for these reads. Also, we enumerate $\theta_{nk}$ using a step size of 10bp in M-step. This is reasonable since the standard deviation of fragment size (30bp) is relative large, which limit the precision we can achieve, i.e. the standard deviation of the estimated pA sites should be larger than that of the fragment (30bp).

We pre-specify the minimum and maximum number of pA sites in the data. For each possible number of pA sites, we perform 5 independent EM inferences with different initialization values for $\boldsymbol{\pi}$, $\boldsymbol{\alpha}$ and $\boldsymbol{\beta}$. Then we choose the best result with the lowest BIC value from the 5 independent inference runs. If the result contains APA components with weight less than 0.01, then we remove these components and use the rest as final results.

In the binning operation, we use 5bp, 10bp, 10bp as the bin sizes of the $\boldsymbol{x}$, $\boldsymbol{l}$, $\boldsymbol{r}$, respectively.

### References

[1] Ivano Legnini, Jonathan Alles, Nikos Karaiskos, Salah Ayoub, and Nikolaus Rajewsky. Flam-seq: full-length mrna sequencing reveals principles of poly(a) tail length control. *Nature Methods*, 16(9):879–886, 2019.
